# Area MT carries acceleration information in a quickly and directly decodable representation

**DOI:** 10.1101/2025.10.14.682245

**Authors:** Penny Shuyi Chen, Alexander C Huk

## Abstract

We sought to better understand the neural representation of visual motion acceleration. A straightforward estimation of acceleration would involve calculating the rate of change of velocity, which itself would be calculated from change in position over time. As it is well-established that neurons in area MT encode velocity, the brain could thus indirectly estimate acceleration by calculating the change in MT’s velocity representations. An alternate mechanism, however, could operate more rapidly and directly. In this case, the brain could exploit interactions between MT’s standard motion encoding and idiosyncratic temporal dynamics of neural responses. Such direct decoding would thus exploit nonlinearities usually ignored in studies of MT coding. We tested between these two theories by measuring from ensembles of MT neurons while two male awake fixating macaques viewed linearly accelerating motion stimuli. Direct decoding of acceleration from MT was possible on faster time scales, and could be done with higher fidelity, than indirect decoding. Distinct motion acceleration information could thus be efficiently read out from rich and heterogeneous MT ensemble responses, regardless of the mechanisms that give rise to various forms of motion tuning that it exhibits. A similar analysis of activity in the medial superior temporal area (MST) did not suggest this later stage of motion processing has a more refined acceleration representation. Together, these results suggest that the brain may opportunistically exploit nonlinear idiosyncrasies of neural responses to efficiently extract behaviorally relevant information on fast time scales, instead of performing explicit calculation of some variables.

**Significance Statement:** This study aims to understand if linear acceleration information of visual motion is extracted and represented in primate visual motion areas MT and MST. By combining large-scale multi-area neuronal recordings, population-level analyses, and a rich set of moving stimuli, we showed that linear acceleration can be decoded from MT activity. Specifically, our results demonstrate that (1) motion acceleration is encoded in MT, and can be directly and quickly decoded from MT ensemble activity, and (2) area MST, despite being a later stage of motion processing, does not refine acceleration representations. These findings call for revisiting how brain areas might efficiently extract behaviorally relevant information from the environment and highlight the importance of temporal dynamics in visual motion processing.

## Introduction

The brains of primates have a specialized pathway primarily focused on the processing of motion and depth information (Andersen1997, .; Czuba et al., 2014; DeAngelis et al., 1998). Area MT is well studied because of tight links between its activity and both motion perception (Born & Bradley, 2005; Newsome & Pare, 1988; Rokers et al., 2009; Rudolph & Pasternak, 1999) and smooth pursuit eye movements (Ilg, 2008; Inaba et al., 2011; Krauzlis, 2004; Thier & Ilg, 2005). Neurons in MT have well-established selectivity for the direction of motion, and also exhibit clear tuning for speed, with columnar and clustered functional organizations further confirming this area’s specialized motion processing (Albright, 1984; DeAngelis & Newsome, 1999; Huk et al., 2002; Liu & Newsome, 2003; Perge et al., 2005). Although these properties have framed MT as a key brain region carrying motion signals relevant to perception and action, it is unclear whether it also encodes information about the rate of change of speed, i.e., acceleration (Lisberger & Movshon, 1999; Price et al., 2005, 2006; Priebe & Lisberger, 2002; Schlack et al., 2007, 2008). In the present study, we sought to build upon prior work to answer this question more definitively.

The degree to which activity in MT (and/or MST, a later stage of motion processing) signals acceleration has been investigated previously. This important prior work has highlighted (and grappled with) complexities related to the necessarily time-varying nature of accelerating stimuli. When a stimulus accelerates, its velocities sweep through a range of values, and hence will stimulate velocity-tuned neurons in a time-varying manner, which complicates standard analyses that typically count spikes in response to a fixed stimulus. Relatedly, time-varying effects of adaptation while viewing time-varying stimuli have further clouded searches for “true” encoding of acceleration that are not explained by adaptation and/or velocity tuning (Lisberger & Movshon, 1999; Price et al., 2005). Finally, simply selecting the right range of speeds and accelerations (and how to sample that range) is difficult, and the values of accelerations used appear to vary widely across prior studies (Calderone & Kaiser, 1989; Cao et al., 2004; Mueller & Timney, 2016; Nakayama & Motoyoshi, 2017; Price et al., 2005; Werkhoven et al., 1992). Thus, the degree to which individual MT neurons encode acceleration remains unclear.

Although the presence and/or quality of single-unit *encoding* of acceleration is not settled, it is more evident that acceleration can be *decoded* from the responses of ensembles of MT neurons (Lisberger & Movshon, 1999; Price et al., 2005; Schlack et al., 2008). The basic idea is that even if MT neurons are only tuned for speed (and not acceleration), their time-varying dynamics could be leveraged by a read-out scheme that figures out what speeds were presented at what times. Work in both anesthetized and awake, fixating macaques has shown that acceleration could be decoded by comparison of fast (transient) time-scale responses to slower (sustained/lowpass) responses (Lisberger & Movshon, 1999; Price et al., 2005; Schlack et al., 2007; Traschütz et al., 2015). Although details of previous decoding approaches differ, they share a reliance on a comparison between what are effectively neural snapshots of velocities at different times. Thus, it remains unclear as to whether acceleration information is directly available from MT ensemble activity, or needs to be calculated from their time-varying responses to time-varying velocities.

We therefore decided to revisit the encoding and decoding of acceleration from MT activity. Although the original study of acceleration decoding by Lisberger and Movshon was hugely influential as an early demonstration of population decoding in vision, and follow up work by Price and colleagues generalized this to awake animals and added refinements to the treatment of temporal dynamics (Lisberger & Movshon, 1999; Price et al., 2005), we decided that these issues deserved investigation using updated tools and analytic perspectives. First, we revisited the question of whether single MT neurons could encode motion acceleration information in a similar way as their encoding of velocity. Second, we decided to simply ask whether MT responses carry information about acceleration, irrespective of the underlying mechanism that might create that tuning. This conceptual shift was motivated by modern emphasis on population decoding, wherein information can be gleaned from neural populations even in the absence of clearly-interpretable and/or explicit tuning at the level of single neurons (Averbeck et al., 2006; Churchland et al., 2012; Quian Quiroga & Panzeri, 2009; Warland et al., 1997). Third, we focused our measurements on a particular set of speeds and accelerations. These were chosen based on perceptually relevant values, and guided by comprehensive assessments of MT speed tuning that mostly occurred after the class studies of acceleration discussed above (Borghuis et al., 2019; Carrasco et al., 2003; De Bruyn & Orban, 1988; Mallery et al., 2010; Nakayama & Motoyoshi, 2017; Nesti et al., 2014). Fourth, we used multi-element electrode arrays to record from multiple MT neurons in each experiment, and applied now-standard population decoders to those simultaneously recorded neural ensembles (Buzsáki, 2004; Saxena & Cunningham, 2019; Stringer et al., 2021; Urai et al., 2022; Vyas et al., 2020).

We therefore investigated acceleration encoding and decoding by presenting a range of linearly accelerating stimuli to awake, fixating macaques while we recorded from ensembles neurons using linear electrode arrays placed into area MT and the medial superior temporal area (MST) (16-80 neurons per session, 648 MT neurons and 461 MST neurons). Consistent with prior work, we found that direction and speed tuning was ubiquitous across MT neurons. But contrary to the conclusions of prior studies, we found that acceleration tuning in MT was quantitatively in the same order as direction and speed tuning. We also found that acceleration could be decoded very well from the ensemble response of MT neurons; the decoder was agnostic to any particular mechanism or physiological property, but could flexibly weight neurons (with heterogeneous dynamics) to best estimate acceleration. Such opportunistic “direct decoding” was not just possible, but was better than decoding velocity and then calculating the derivative (“indirect decoding”). Direct acceleration decoding was thus possible with low latency and across short time scales, suggesting it is of advantageous behavioral relevance. Finally, a similar analysis of MST neurons did not suggest that this area reflected a later, more explicit or more refined stage of acceleration representation. Our work thus confirms the capabilities of population decoding to identify variables of interest that may not be obviously and explicitly encoded at the single neuron level, and shows that idiosyncratic and heterogeneous response dynamics can be opportunistically used to exploit nonlinear selectivities that are inscrutable at the single unit level but which are information-rich at the population level.

## Materials and Methods

### Experimental design

#### Stimuli

Visual stimuli were processed and presented through Psychophysics Toolbox (Kleiner, 2007) in MATLAB 2020b (Mathworks) with integration of DATAPixx2 video I/O hub (VPixx Inc.) for precise timing (Eastman & Huk, 2012). Both monkeys (Monkey K and Monkey L) were chaired at the recording rig with a viewing distance of 101 cm. Visual stimuli were projected onto a 182.22 cm by 102.5 cm polarization-preserving rear-projection screen by an PROPixx 3D projector (VPixx Technologies) running at 120Hz frame rate with pixel resolution at 1920 x 1080 through a DepthQ 3D polarization module, a high-speed circular LCD polarizer for passive interleaved-stereo stimulation. This setup allows to cover 84.11 x 53.81 degree of visual angle with 20.071 pixel per degree and < 1% crosstalk between the eyes. Binocular eye positions and movements were tracked by Eyelink1000 (SR Research) and sampled at 1kHz. Fluid reward was delivered by a computer-controlled precise programmable syringe pump (New Era Pump Systems, Inc.) at the end of each trial. Within each experimental session, we first functionally mapped the neurons recorded by the array to decide on the preferred-null direction axis, as well as receptive field size and location, which we then used to generate the acceleration stimuli. Full details of our mapping battery (Fig 1a - MT example session functional mapping results), which also included tuning measurements for a variety of motion and depth variables, are described in the *Electrophysiology* section later.

**Figure 1.**
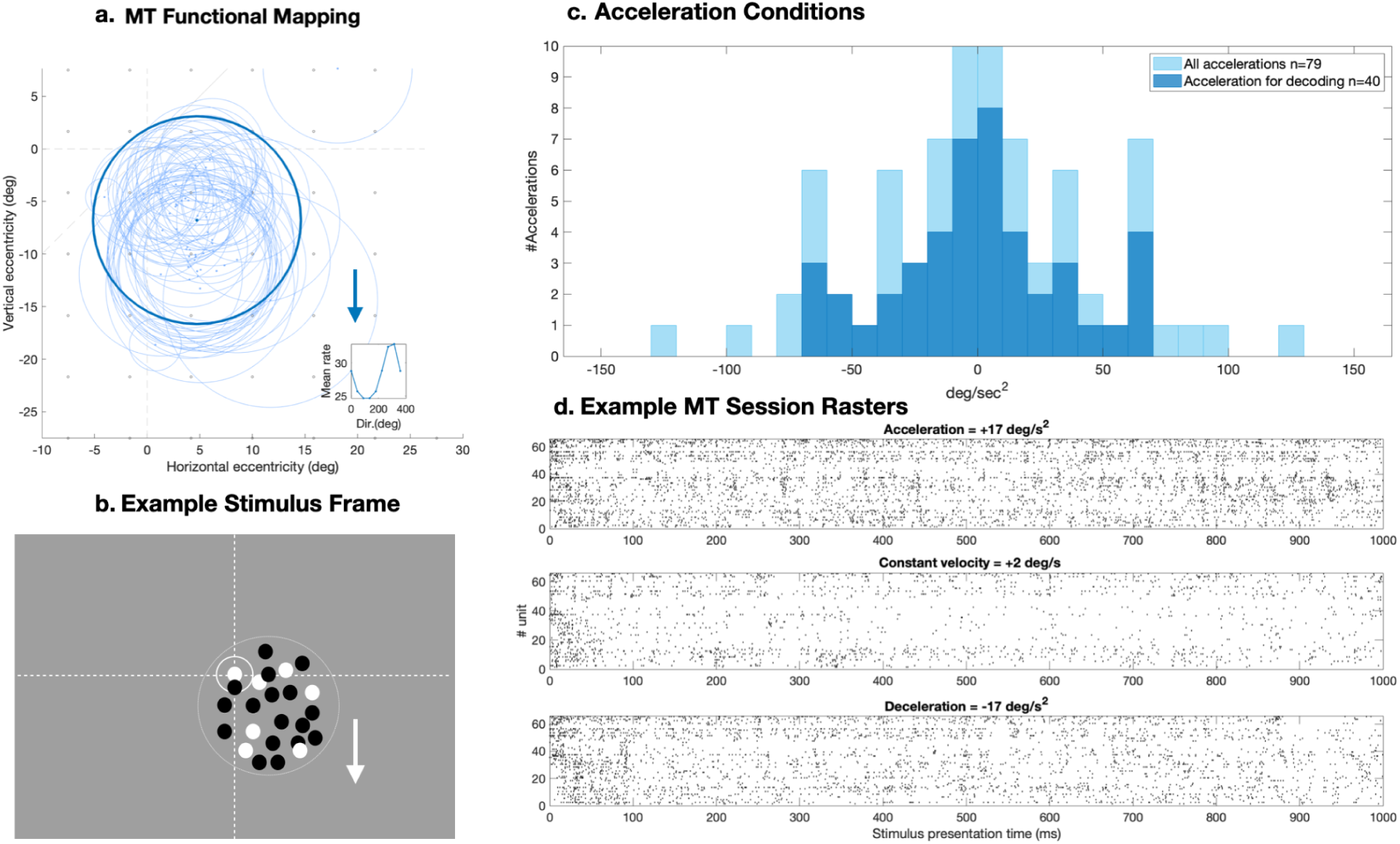
Example stimulus and experimental design. **a**, MT functional mapping: light blue contours show individual receptive fields (RFs) and the bold contour indicates the ensemble RF which determines the location and size for the acceleration stimulus envelope in b. The blue arrow shows the direction of preferred-null motion axis (preferred direction at 270 degrees) of the recorded population with the inset plot showing the population direction tuning. **b**, Example stimulus: The bull-eye circle represents the fixation dot shown throughout the trial. The large white circle, which was not presented in the actual experiment, indicates the size of the motion dots field envelope. The arrow represents the positive acceleration direction (preferred motion direction at 270 degrees) determined by MT functional mapping in a. **c**, Histogram of acceleration conditions: solid color histograms show 40 unique acceleration conditions that were used for decoding analysis. The transparent color histograms show the full 79 unique acceleration conditions derived from the full condition matrix. **d**, Example MT ensemble rasters (Monkey K): top–example acceleration condition, middle–example constant velocity condition, bottom–example deceleration condition.

For the main linear acceleration motion experiment, subjects were required to maintain fixation on a central point for 250 ms, after which moving dot stimuli were presented. The scheme of the motion dots stimulus frame is shown in Figure 1b. In each trial, two 1s long linear acceleration motion dot field stimuli were presented sequentially. The moving dot field consisted of a circular patch of light and dark anti-aliased dots moving coherently and frontoparallel to the observer. We pseudorandomly selected stimuli from a design matrix that contained all possible combinations of 15 possible starting velocities ( -64, -32, -16, -8, -4, -2, -1, 0 , 1, 2, 4, 8, 16, 32, 64 deg/sec) and 15 possible end velocities (identical to starting velocities set), with positive values indicating motion in the preferred direction and negative values indicating null-direction motion. Thus, there were 255 start-end velocity combinations that yielded 79 unique acceleration conditions (+/- deg/s^2^) covering a large sample space of motion velocities and linear accelerations. For all analyses in this paper, we restricted our analyses to (a) completed trials in which the acceleration or deceleration were in the preferred direction of the neurons under study, and (b) trials in which the direction of motion did not reverse (a byproduct of our fully-crossed design of starting and ending velocities that could be in either the preferred or null direction). We incorporated equal amounts of preferred and null direction speeds in order to limit long term direction-selective adaptation, but plan to consider the null and reversing-direction conditions separately (Calderone & Kaiser, 1989). This leaves 40 unique acceleration (in the preferred direction) conditions, derived from the full condition matrix, which were used for decoding analysis (Figure 1c - histograms of unique acceleration conditions). We also note that for analyses of encoding and (direct) decoding of acceleration, we marginalized over (i.e., ignored) the starting velocity. For encoding analyses at both single cell and ensemble levels, the full set of acceleration conditions are used for computing acceleration tuning curves.

Monkey K’s example session ensemble rasters of MT activity during 3 example trials are shown in Figure 1d: one acceleration trial, one constant velocity trial, and one deceleration trial. A trial was aborted if the animal broke fixation during either of the motion stimulus presentations. There was a 250 ms inter-trial-interval during which we administered fluid reward for trial completion. Each experimental session acquired 17 to 20 repeats of all conditions.

#### Electrophysiology

We performed electrophysiological recordings from area MT and MST of two male rhesus macaque monkeys (Monkey K and Monkey L, 20 and 11 years old). The extracellular recordings were performed with Plexon 32 channel linear electrode arrays (U-probe and S-probe), with 50um within and 100um between stereo-pair site spacing that gives a 1.5mm total span. A titanium recording chamber and 3D printed recording location grid (19mm x 25mm with 1mm spacing) customized to each individual animal was placed over the superior temporal sulcus (STS) and intraparietal sulcus (IPS) to allow linear array penetrations tangential to the cortical surface at MT. This design gave us the opportunity to record from columns of MT or MST neurons for each session (which yielded rather similar tuning across the ensemble within each experiment session), as well as to sample different columns across sessions. The chamber placement for each animal was guided by macaque average stereotaxic coordinates (Paxinos et al., 2003) and each animal’s individual structural MRI scan and cranial landmarks.

For each recording session, MT or MST was first identified using the recording location coordinates on the recording grid and electrode penetration depths, and then verified by functional mapping. The functional mapping consisted of two parts: 1) identifying the individual and aggregated spatial receptive field locations across the recorded units, and 2) concurrent mapping of 2D motion direction and binocular disparity tuning. First, while the subject was fixating at the center of the screen, a hand-mapping stimulus consisting of a circular field of moving dots with on-fly experimenter control to stimulus properties (e.g. motion dots field location, size, speed and density) was presented to roughly map the visual responsive area across the recorded units (i.e. coarse population receptive field). Then a 7x7 motion dots field grid was placed over the center of coarse population RF to determine individual unit’s spatial RF location. Rapid random sequential presentations of 3-5 deg diameter motion dots field with uniform density were shown during stable fixations. Because we were recording with an orthogonal approach to the cortical sheet (and thus as would be expected from recording a column of neurons), the individual RFs were generally clustered and spiraled together in space (Figure 1a, light blue contours). The diameter of the uniform field of moving dots for motion direction-disparity tuning was scaled to span the aggregated RF (15-30 deg).

For determining recorded units’ classic motion tuning, 2D direction and binocular disparity tuning were measured concurrently with 500ms presentations of a fronto-parallel plane of moving dots with randomly drawn 2D motion direction (0-360 deg with 45 deg increment) and disparity offset (9 steps that spans +/-1.5 deg of horizontal disparity) at a speed of 10 deg/sec. In each session, the motion direction that maximally drove the highest number of recorded units was chosen as the population’s preferred (2D) motion direction and thus determined the positive acceleration direction. Together, the results of this functional mapping were used to define the dot field’s location, diameter and preferred-null motion direction axis of acceleration for the linear acceleration stimuli (Figure 1b).

A total of 20 recording sessions in MT (Monkey K 11 sessions, Monkey L 9 sessions) and 11 sessions in MST (Monkey K 9 sessions; Monkey L 2 sessions) were performed. Spike sorting was done in KiloSort 2.5 (Pachitariu et al., n.d.) followed by manual curation in Phy2. Only MT and MST units that were labeled as ‘single unit’ (the ‘Good’ group after manual curation) were included in the analysis. A total of 648 MT units (Monkey K 430 units, Monkey L 218 units) and 461 MST units (Monkey K 415 units, Monkey L 46 units) were identified and included in the analysis.

### Statistical analysis

#### Variations of Poisson Independent Decoders for estimating acceleration

To perform single-trial decoding of acceleration from population neuronal responses, we adapted a linear parametric decoder, the Poisson independent decoder (hereafter, “decoder”), which was previously applied to decode the activity of neuronal populations (e.g., Bonnen et al., 2020; Graf et al., 2011). We used a variety of decoders based on this standard architecture that vary along two dimensions. First, we employed either a “direct” decoding scheme (based on measured acceleration tuning), or an “indirect” scheme (based on measured velocity tuning, which then required an additional step to calculate acceleration from the rate of change of the decoded velocities). General schematics of the direct-acceleration decoder and the indirect-velocity-based decoder are shown in Figure 2.

**Figure 2.**
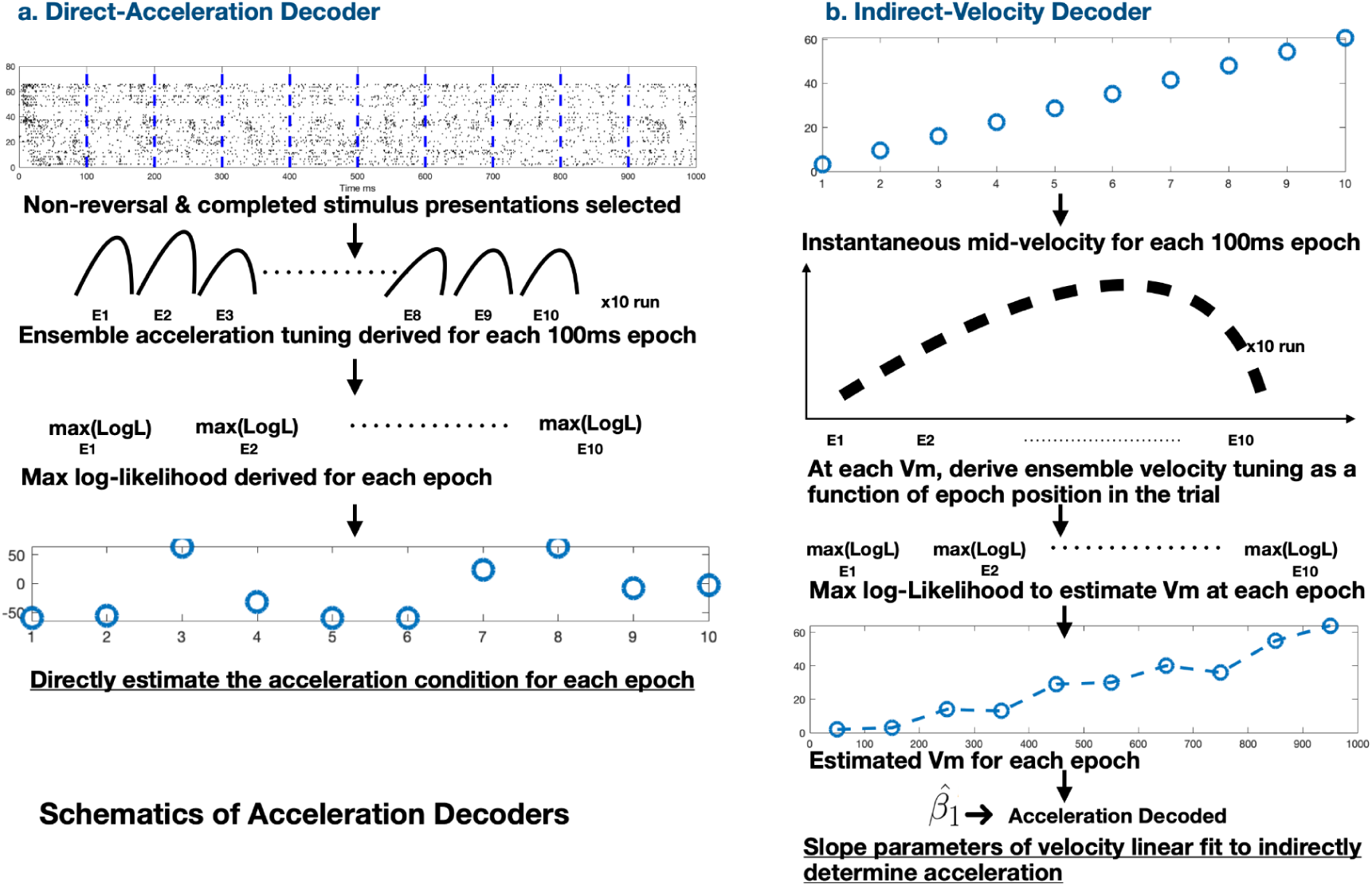
Acceleration Decoder Schematics. a, Schematics of dynamic direct-acceleration decoder and b, Schematics of dynamic indirect-velocity-based decoder.

Second, due to the time-varying nature of acceleration, we employed either “dynamic” or “static” decoders. To generate ensemble acceleration or velocity tuning for dynamic decoders, each one-second long stimulus presentation was divided into ten 100 ms epochs, with the first epoch generally capturing the onset transient period, the middle 8 epochs encompassing the sustained period of response to the stimulus, and the last epoch capturing the offset-transient period. Tuning was calculated for each 100 ms epoch independently as the pooling weights, assuming the neuronal tuning is not fixed but has been rapidly computed with its own temporal dynamics over the stimulus presentation and thus affects the decoding strength dynamically. In contrast to the dynamic decoders, the static decoders decoders used tuning functions measured either from onset transient response only (i.e. the response to the first 100ms of stimulus presentation) or during the full stimulus presentation, and applies the same tuning weights for every 100ms epoch for decoding-the same way that it has been applied to simpler cases like orientation. These ‘static’ decoders assume the full temporal dynamics of visual motion encoding is not necessarily needed to efficiently and accurately compute or estimate ongoing motion’s acceleration-either a snapshot (i.e. transient-tuning) or a summary (i.e. full-tuning) of the visual motion is sufficient.

Specifically, there are two variations of static decoders: 1) Transient-tuning decoder, and 2) Full-tuning decoder. The transient decoder suggests that the onset-transient period tuning can provide sufficient information about the ongoing motion acceleration in a rather short amount of time. Therefore, this information is carried on throughout the stimulus presentation, and the decoder’s pooling weights are fixed to be the ensemble transient period tuning. On the other hand, the full-tuning decoder considers the population neuronal tuning that is computed from the full stimulus presentation that summarizes the neuronal dynamics from motion onset to offset is essential to precisely decode motion acceleration. Hence, the pooling weights are fixed to be the neuronal tuning calculated with spike counts of the entire stimulus presentation.

In summary, the pooling weights for our acceleration and velocity-based decoders are determined by the choice and assumption of motion tuning dynamics.

The inference from the decoder is represented by the log-likelihood function, which is the conditional probability of observing the unit responses given a stimulus, evaluated across the chosen stimuli (Graf et al., 2011; Jazayeri & Movshon, 2006; Ma et al., 2006). For our various decoders, the likelihood function linearly combines neuronal responses across units using the pooling weights derived from their motion tuning functions, as follows:

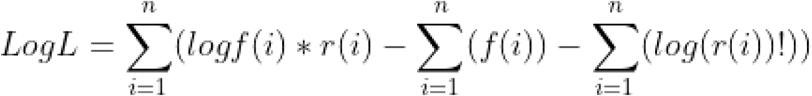

where f is the ensemble tuning, and f(i) represents the unit motion tuning (acceleration or velocity tuning) at a specific acceleration or velocity for a specific time epoch; *r* is the population response, in which *ri* represents the unit response for a given stimulus.

For each session, we trained and tested the decoders with non-reversal stimulus presentations, in which there was no change of dots motion direction, only. 90% of the selected stimulus presentations were used for training to derive ensemble tuning and the rest were used as the testing set to generate log-likelihood functions to dynamically estimate the acceleration or velocity of the stimulus at each 100ms epoch. Each decoder was run for 10 times and the maximum values of the averaged log-likelihood functions were taken as the decoder estimates. These estimates then will directly (direct-acceleration decoders) or indirectly (velocity-based decoders) determine the decoded acceleration condition of the given stimulus. Further description about direct-acceleration decoders and velocity-based decoders and their variations is described in the Result section.

#### Decoding quality analysis

For each recording session, mean adjusted-R-squared values against the line of unity with bootstrapping (10000 iterations with replacement on the testing stimulus presentation set) and 95% confidence intervals were computed. The mean adjusted-R-square value demonstrates the decoding quality for the entire stimulus presentation, the decoding quality without the onset-transient period, or the decoding quality at each epoch to show the dynamic of acceleration decoding performance across stimulus presentation. (See Figure 4 for Monkey K’s example MT session’s decoding results from two decoders.)

#### MT/MST motion tuning indices

To characterize basic neural selectivity and to investigate encoding of acceleration and related variables that MT is known to represent, we calculated several tuning indices in the similar way to previous studies of MT (Liu & Newsome, 2003; Price et al., 2005): 1) direction tuning, 2) velocity tuning , 3) acceleration tuning and 4) binocular disparity tuning. Tuning functions for velocity, acceleration, disparity were fitted with a smoothing cubic spline function (to avoid making parametric form assumptions, which we realized might be incorrect especially for acceleration), while the responses for direction tuning were fitted with the well-established circular gaussian (Von Mises) function (Liu, Newsome 2003). After extracting the fitted parameters, these four tuning indices are calculated with the same basic equation:

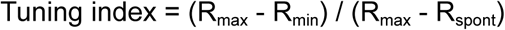

where R_max_ and R_min_ represent the highest and lowest responses of the tuning curve and R_spont_ as the spontaneous activity. A value of 0 indicates no tuning to that stimulus dimension, and a value >1 indicates the unit or population is excited by its preferred stimulus but suppressed by other non-preferred motion stimulus in the evaluated motion dimension.

For comparison to prior work that has studied MT’s encoding of acceleration, we also calculated an acceleration-deceleration index using the formula from Price et al, 2005:

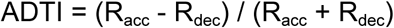

in which R_acc_ is the max response of the accelerating conditions of the tuning curve and Rdec is the max response of the decelerating conditions of the tuning curve.

The spike counts that are used to compute direction and binocular disparity tuning were taken from our functional mapping battery (see Electrophysiology section). The velocity tuning index was calculated from the stimulus presentations in the main experiment that had no acceleration (i.e., constant motion velocity). The acceleration tuning index was computed with the spike counts from all available acceleration experiment stimulus presentations collapsed over start-end velocity combinations.

#### Receptive field parameters

MT and MST’s receptive fields were fitted with a 2D Gaussian. Adjusted-R-squared values were computed for each unit’s RF estimate as the measure of individual RF fitting quality. The units with a value greater than 0.9 were included into the analysis.

For example units and sessions’ figures/visualizations, the same session and unit (from the selected session) data are used (Monkey K: session 20220413, unit 63. Monkey L: session 20220622, unit 36).

#### Code Accessibility

The code for main acceleration experiment, acceleration and velocity based decoder and analysis with example session data are available on GitHub repository: https://github.com/PennyShuyiChen/MTMSTAccelerationEnDeCoding. Example code and stimulus presentation for RF mapping and MT/MST functional mapping are available at https://www.visualstimul.us/code.html (Czuba, 2023).

## Results

We first measured the basic tuning properties (i.e., direction, velocity and binocular disparity tuning) and receptive field locations of the sampled MT and MST neurons. We then measured the time course of MT and MST ensemble responses to linear acceleration motion stimuli with various starting velocities in two fixation-trained male rhesus macaques. Our main goals were threefold: (1) to test whether individual MT neurons exhibit explicit tuning to acceleration; (2) to evaluate whether direct decoding of acceleration is possible from MT responses and how the performance of such a decoder compares to indirect estimation of acceleration based on changes in decoded velocity; and (3) to assess whether the representation of acceleration information in area MST is more explicit and/or of higher fidelity than that in MT. We found that, within the range of velocities and accelerations we tested, MT neurons exhibited explicit tuning for acceleration. This likely contributed to the fact that direct decoding of acceleration was possible from MT ensembles, and that it was better and faster than decoding velocity and then estimating acceleration from that. Finally, responses in MST did not suggest a more refined representation of acceleration. Taken together, these results reveal tuning in MT that may have been overlooked given prior experimental approaches, and suggest that acceleration information could be “read out” from MT populations at behaviorally-relevant timescales.

### Area MT neurons exhibit tuning to acceleration that is quantitatively similar to other motion tuning properties

We first tested whether individual MT neurons exhibit explicit tuning to visual motion acceleration in a way similar to their encoding of velocity. This was motivated by prior studies that did not unambiguously answer this question, and which suggested that any modulations seen in MT responses as a function of acceleration might be “explained away” by tuning to velocity and interactions with temporal dynamics - such as transient/sustained phases of responses, and/or adaptation. To address this issue, we measured single unit tuning curves for acceleration, quantified them using a simple discriminability index, and then compared that tuning strength to the same metrics estimated for direction, velocity - stimulus features that have been studied more extensively in MT, and for which there is little controversy about the presence of tuning.

The reason for an ambiguous state of understanding despite a seemingly simple question is due to two primary factors. First is the issue of stimuli and conditions. We focused on a relatively limited range of speeds and accelerations, finely sampling acceleration values within a perceptually-relevant range (Lappin et al., 2009; Naseri & Grant, 2012; Priebe & Lisberger, 2004; Schmerler, 1976; Traschütz et al., 2015; Watamaniuk & Heinen, 2003).

Furthermore, we note that the same acceleration could be generated with different starting velocities. We grouped trials by the acceleration, which could mean that some conditions had trials that had different starting and ending velocities, but which had the same acceleration. The second issue is related to this last point, but is more conceptual. Our perspective is that, if the observer simply wants to extract acceleration information from a neural population, the mechanistic source of that information is not important to the decoding process. Thus, we did not attempt to “explain away” any acceleration tuning, and in fact were interested to see if the brain might opportunistically exploit interactions between velocity tuning and temporal nonlinearities in response.

Example single unit acceleration tuning curves (Figure 3a bottom panels) show clear changes in neural response as a function of acceleration. Although the shape of these acceleration tuning curves does not follow a well-established functional form, they have a clearly systematic shape. One example neuron increases its responses to positive accelerations (i.e., increasing speed in the preferred direction) which then falls off gradually at higher acceleration rates (Monkey L, 3a bottom left panel); it is slightly suppressed by decelerations and that suppression similarly decreases for higher values. The other example neuron decreases its responses to decelerations (i.e., decreasing speed in the preferred direction) which then falls off gradually at higher deceleration rates; the cell is slightly facilitated by accelerations (Monkey K, 3a bottom right panel). Although quantitatively different, the general form of acceleration tuning functions is similar in these two example neurons.

**Figure 3.**
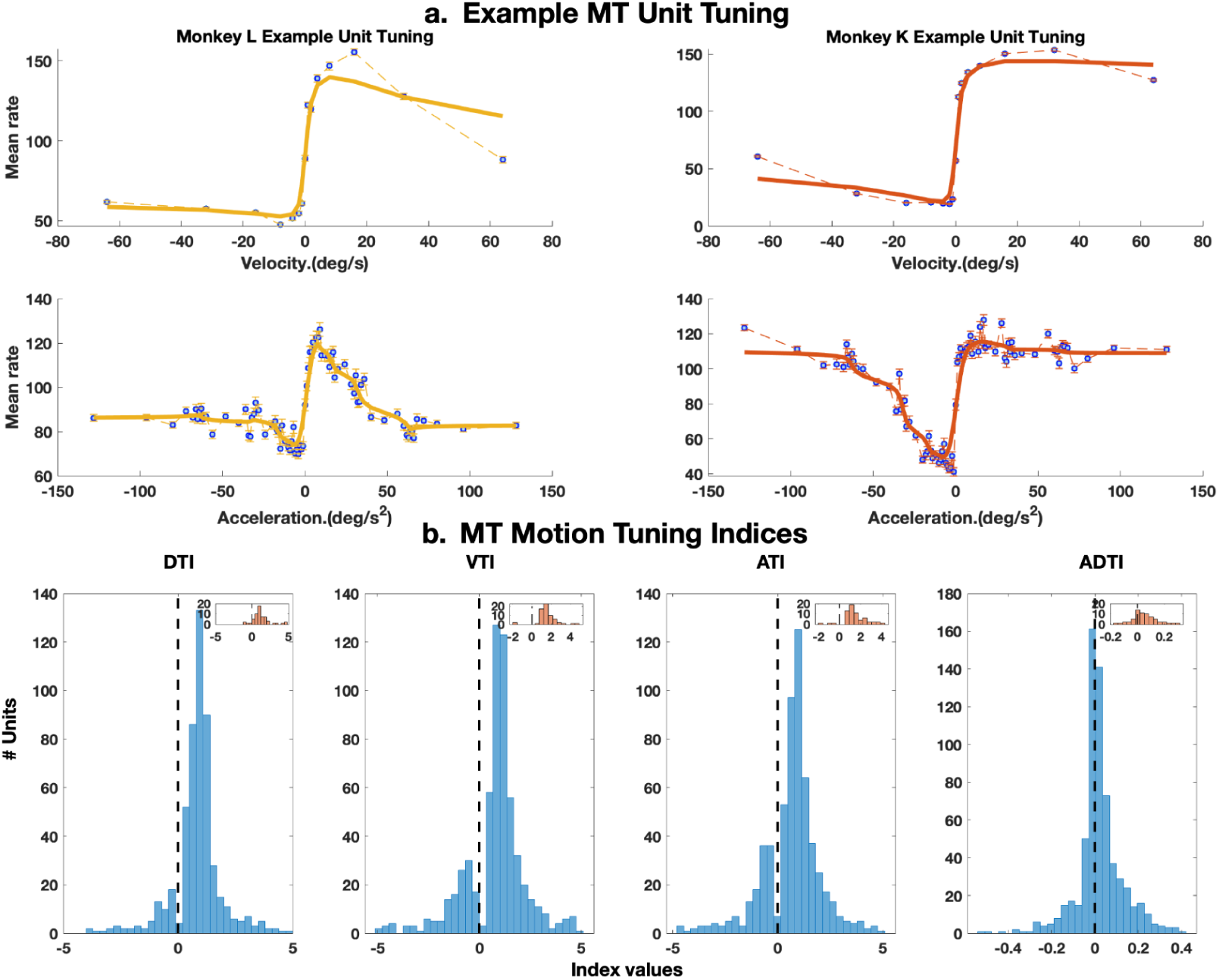
MT motion tuning. a, Example MT unit velocity and acceleration tuning curves. Top–bottom: velocity tuning–acceleration tuning, left–right: example unit from Monkey L -Monkey K. Error bars reflect +/- 1SEM with the dashed lines connecting the actual mean rates at each velocity or acceleration measured, and the bold lines representing the Gaussian fits. b, Histograms of Motion Tuning Indices. Left–right: Direction tuning index, Velocity tuning index, Acceleration tuning index, and acceleration-deceleration tuning index. The insets showed the MTI distributions of a single example MT session from Monkey K. (The example Monkey K unit in a is pulled from the example ensemble of b insets).

Interestingly, the velocity tuning for these example neurons (shown above the acceleration tuning curves, Figure 3a top panels) follow a rather similar pattern. These tuning curves are directional, with facilitation in the preferred direction, suppression in the null direction, and cell-to-cell variations in how strong the facilitation and suppression are. To perform a fair quantitative comparison between velocity and acceleration tuning, we fit a cubic spline to the measured data points, and calculated a simple and classic nonparametric tuning index that coarsely captures how strongly modulated the cell is by changes in acceleration or velocity (see Methods).

At the population level, we found that acceleration and velocity tuning indices were very similar (Figure 3b, panel 2 velocity tuning and 3 acceleration tuning: t(647) = 1.449, *p* = 0.148, Velocity tuning: M = 0.776, SD = 1.470 with 95% CI [0.659, 0.894], Acceleration tuning: M = 0.658, SD = 1.362 with 95% CI [0.258, 1.069). This suggests that acceleration tuning in MT is not just explicit, but of approximately the same quality as for that of velocity (AVI and VTI distributions are not significant from each other: t(647) = 1.4486, *p*=0.148). We also used the same approach to quantify direction tuning and binocular disparity tuning, and again found similar degrees of tuning (Direction tuning: M = 0.857, SD = 0,950; binocular disparity tuning: M = 0.950, SD =1.22. t(647) = -1.295, *p* = 0.196). The similarity here suggests that, at least at the coarse level of a tuning index, acceleration is on par with more classical tuning dimensions in MT.

Specifically (Figure 3b), 74.54% (n = 483) of the recorded MT units had preferred motion acceleration with 38.12% (n = 247) excited by their preferred acceleration and inhibited by other accelerations, and the remaining 25.46% (n = 165) were inhibited by all accelerations. The recorded MT neurons from Monkey K and L showed similar acceleration tuning strength. MT single unit’s velocity tuning is analogous to acceleration tuning: 77.01% (n = 499) of the neurons had velocity preference, and 49.85% (n= 323) were excited by their preferred velocity but inhibited by other velocities, but 22.99% (n = 149) of the neurons were inhibited by any velocities. Furthermore, direction and binocular disparity tuning indices also demonstrated comparable distributions with 85.92% (n = 556) of the neurons showing direction and disparity preference, 44.97% (n = 291) and 49.54% (n = 321) being excited by the preferred stimuli, but 14.08% (n = 92) were inhibited by all motion directions and disparities. As shown in Figure 3b, all MTI distributions are different from 0 (DTI: M=0.857, SD=1.070, t(647)=18.175, *p*<0.001; VTI: M=0.776, SD=1.470, t(647)=13.010, *p*<0.001; ATI: M=0.658, SD=1.362, t(647)=11.854, *p*<0.001;).

Finally, we also calculated the acceleration-deceleration tuning index for each unit to first compare to the measurements by a previous study that used the same metric (Price et al., 2005), and also to more directly test whether MT neurons prefer fixed velocities or accelerations. Overall, similar to the previous study’s results, the two animals exhibited similar distributions of acceleration-deceleration tuning index values, demonstrating a preference of positive acceleration across the MT population. In detail, 58.49% (n = 379) of the MT neurons preferred accelerations over decelerations, and 36.57% (n = 237) preferred decelerations over accelerations. Only a small proportion of recorded MT neurons preferred constant velocities over accelerations or decelerations (4.94%, n = 32) (Figure 3b Acceleration-Deceleration tuning index distribution is significantly different from 0, ADTI: M= 0.02, SD = 0.11 with 95% CI [0.013, 0.030], t(647)=5.050, *p*<0.001).

To summarize, the tuning of single units in MT is comparable across motion dimensions, and acceleration tuning is not notably different from that seen for more established measures of direction and speed (as well as disparity). The presence of comparable acceleration and velocity tuning at the single unit level thus re-opens the issue of whether acceleration is inferred from MT based on the decoding of velocity and then performing an additional computational step to estimate the rate of change of decoded velocity, likely in a processing stage beyond MT, or whether acceleration might be in fact be directly decoded from MT.

### Area MT carries acceleration information for efficient and direct decoding

To investigate if the motion acceleration information could be directly extracted from ensemble MT activity, we evaluated the performance of several types of decoder. In all of these decoders, the same basic principles applied, as has been done in other work that used poisson independent decoders for orientation and direction (Bonnen et al., 2020; Ghanbari et al., 2019; Graf et al., 2011; Jazayeri & Movshon, 2006). Here, the primary two types of decoder are (a) a “direct acceleration” decoder, which takes the acceleration tuning curves measured for each neuron to generate ensemble tuning, and then fits decoding weights to estimate the acceleration presented on each trial; and (b) a “velocity-based” decoder, which takes the velocity tuning curves of each neuron, fits decoding weights, and then estimates the velocity over time. This velocity-based decoder thus requires a second step, which is calculating the rate of change of the decoded velocities to arrive at a decoded acceleration. In this exercise, we assume that this extra step (i.e., taking the time derivative of velocity estimates) is performed perfectly, and thus do not incorporate additional noise in this second stage. The assumption of a noise-free extra processing step proves not to be problematic given the pattern of results we soon describe. (See Figure 2 for general schematics of the Direct-acceleration and Indirect / velocity-based decoders.)

Because acceleration is a time-varying dimension, we computed either the ensemble acceleration or velocity tuning curve for each 100ms epoch, and then used those tuning curves to estimate acceleration or velocity as a function of time. For the direct-acceleration decoders, we estimated each epoch’s acceleration by taking the mean of those values obtained from 10 runs. Whereas velocity-based decoders first decoded velocity for each 100ms epoch. Then we estimated acceleration by estimating the rate of change between pairs of estimated velocities of the adjacent epochs (i.e., equivalent to fitting a line to pairs of velocity estimates and then taking the slopes of the fitted lines).

Figure 4 shows the overall performance of these two decoders as applied to a same experimental session in which we simultaneously recorded from 80 MT neurons of Monkey K. To determine a full trial’s estimated acceleration, we took the average of the estimates of each epoch from direct-acceleration decoder or epoch pairs from velocity-based decoder (Figure 4 a, Direct-acceleration decoder; and b, indirect-velocity-based decoder, first column). The direct-acceleration decoder is clearly good, with decoded accelerations matching actual (presented) accelerations well, evidenced by a cloud of points that cluster around the line of unity. The velocity-based decoder is good but not quite as good, with decoded accelerations also following the line of unity, just less tightly than the direct-acceleration decoder. Interestingly, as we took the average of the estimates of the sustained response epochs (i.e., epoch 2-10) and left out the onset-transient epoch (i.e., the first 100ms epoch) for the decoders, the decoding performance decreased as the points spread more away from the unity line (Figure 4 a&b, second column). These visual impressions are confirmed via quantification, where we calculated the goodness of fit of the line of unity in accounting for the decoding points. To illustrate the decoding performance at different timescales, the adjusted-R^2^ between decoded accelerations and actual accelerations for each trial were computed with 10000 iterations of bootstrapping with 95% confidence intervals.

**Figure 4.**
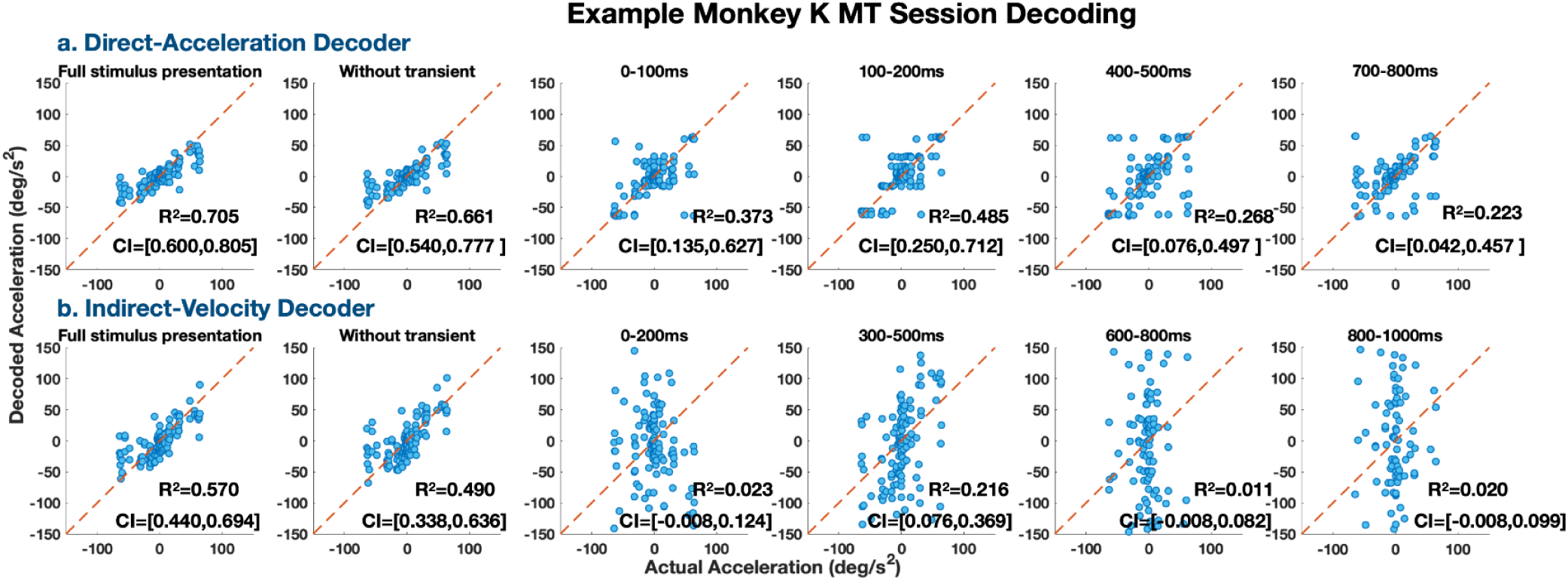
Monkey K example session time course of acceleration- and velocity-based decoding. Monkey K’s MT example session decoding results from **a.** dynamic direct-acceleration decoder and **b.** dynamic velocity-based decoder. Left–right: 1) decoded acceleration based on estimates of response to full-stimulus presentations, 2) decoded acceleration based on estimates of response to stimulus presentations without the onset-transient epoch, and 3-6) decoded acceleration based on estimates for 4 example epochs.

Notably, the difference in performance of the decoders is more striking when one inspects decoding performance in finer time bins. Figure 4 a&b columns 3-6 show the decoding performance of these two main decoders for example time epochs. The direct-acceleration decoder performs similarly across all time bins, with decoding performance less good than the full-trial average decoding, but no striking categorical differences over time or in comparison to the full-trial performance. In contrast, the velocity-based decoder suffers at this fine time resolution, as the decoding scatter plots reveal very broad scatter, consistent with decoded values that have very little relation to the presented values. This is especially evident when one notes that for zero acceleration, there is a vertical streak of decoded points spanning almost all possible values. Thus, the direct-acceleration decoder appears fairly robust at fine time scales, while the velocity-based decoder appears to need long time scales to produce accurate estimates of acceleration.

### Time course of acceleration decoding suggests a dynamic read-out and a distinction between transient and sustained phases

Although the distinction in fine time-scale decoding performance between direct-acceleration decoding and velocity-based decoding already suggests that direct encoding/decoding of acceleration might be valuable for behaviorally-relevant time scales, we further investigated time-varying versions of these decoders to better understand this striking distinction. Specifically we used (1) dynamic, 2) transient, and 3) full-tuning versions of the two decoders. In order to reveal the decoding efficiency and dynamics, as each trial was divided into ten 100ms epochs, the first epoch includes the motion onset transient period’s activity and the rest captures the more sustained activity at different time points. The selection of the decoder pooling weights depended on which variation of the decoder was running (see Method section-

#### Variations of Poisson Independent Decoders for estimating acceleration for details)

In short, dynamic decoders apply a different tuning weight calculated specifically for each epoch, transient decoders take the transient period’s acceleration or velocity tuning (from first epoch only) as the weight for all epochs and full-tuning decoder use the acceleration or velocity tuning calculated based on full-trial firing rates as the weight for every epoch. Therefore, compared to the dynamic decoders, the transient and full-tuning decoders are essentially the ‘static’ acceleration decoders.

We first qualitatively looked at the acceleration tuning derived from the dynamic-acceleration decoder at single cell and ensemble level (Figure 5, a, Example single cell tuning; b, Example ensemble tuning). In general, at the single cell level, the acceleration tuning for sustained epochs are qualitatively similar to each other (epochs of 100-1000ms), but there are also clear quantitative differences: the first epoch exhibits notably higher overall firing rates, and the precise form of the tuning curve continues to change over time (with the larger changes occurring in the first half second, with more modest changes occurring in the second half second). This visual pattern is recapitulated in the ensemble level tuning. Altogether, this brief investigation of dynamic acceleration tuning makes it clear that distinct information might be gleaned from dynamic decoders that take this temporal nonlinearity into account.

**Figure 5.**
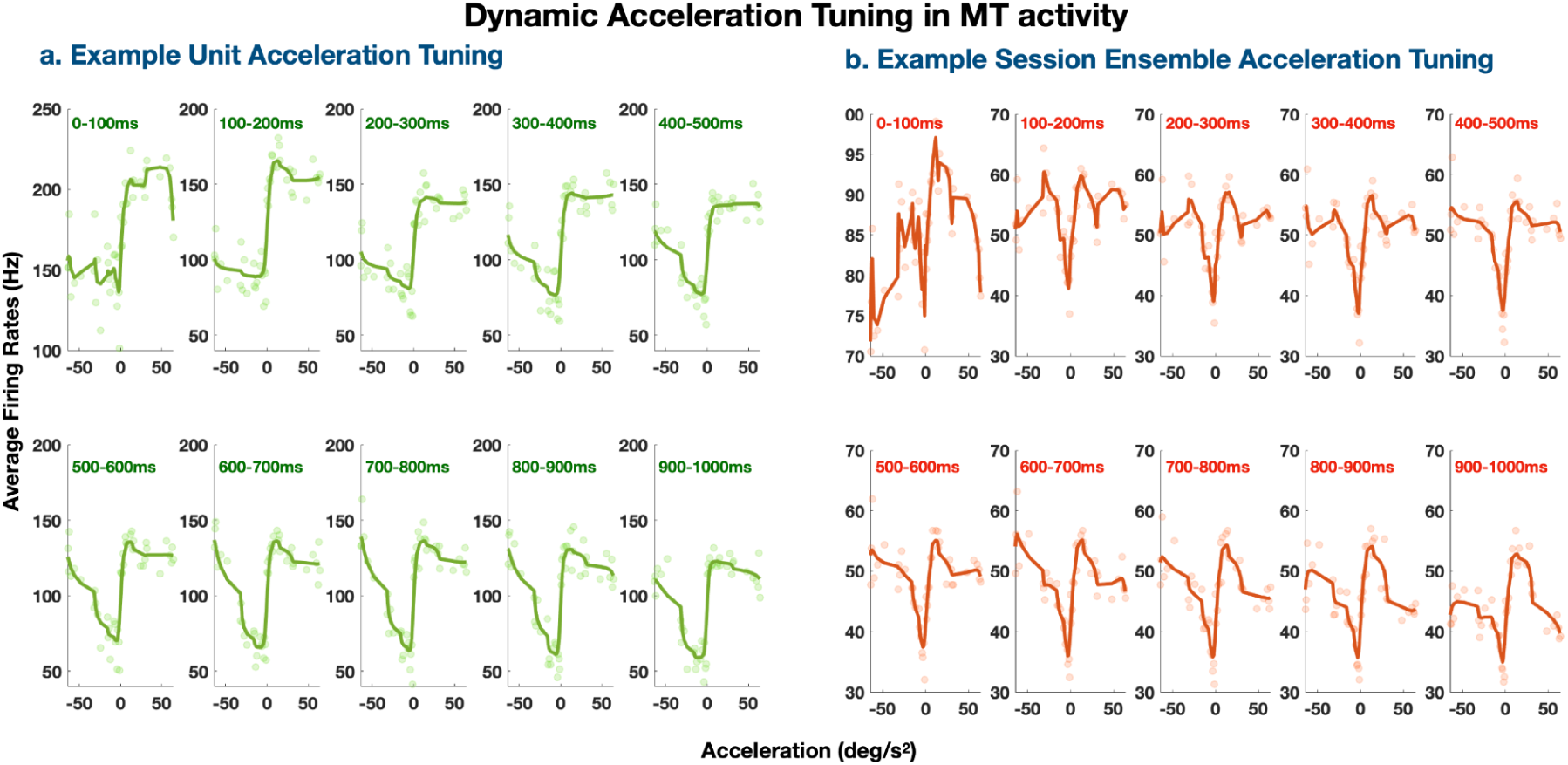
Acceleration tuning dynamics across the stimulus presentation. **a**, Example unit acceleration tuning: Single cell level acceleration tuning curves derived for each 100ms epoch from the same example MT unit of Monkey K’s example session. Although there is a general qualitative similarity across epochs, the tuning curve changes quantitatively over the first few 100 msec (top row), finally settling down during the second half of the stimulus (bottom row). **b**, Example session ensemble acceleration tuning: Population level acceleration tuning curves derived for each 100ms epoch from the same Monkey K’s example session. Transparent circles represent the actual tuning data point at a specific acceleration condition with Gaussian fit of the data overlaid (solid colored lines). Although the precise shape of a population-averaged tuning curve is not straightforward to interpret and is visually cryptic, it is clear that even the average tuning over neurons is dynamic, with the largest changes occurring over the first few 100 msec.

To better summarize and compare direct-acceleration decoding to velocity-based decoding performance and understand the temporal dynamics of MT extracting motion acceleration, we derived average adjusted-R^2^ between decoded acceleration and actual acceleration for all three time-varying versions of the decoders across all MT recording sessions from both monkeys (Figure 6 for dynamic decoders’ results and Figure 7 for transient and full-tuning decoders’ results). Overall, two monkeys had similar decoding performance to each other’s for all variations of the two decoders (e.g. Dynamic acceleration tuning, Monkey K and L: t(200.172), p = 0.876, M = [0.318, 0.313], SD = [0.231, 0.211]; Dynamic velocity decoder: t(176.760) = -0.584, p = 0.560, M = [0.093, 0.102], SD = [0.114, 0.108].)

**Figure 6.**
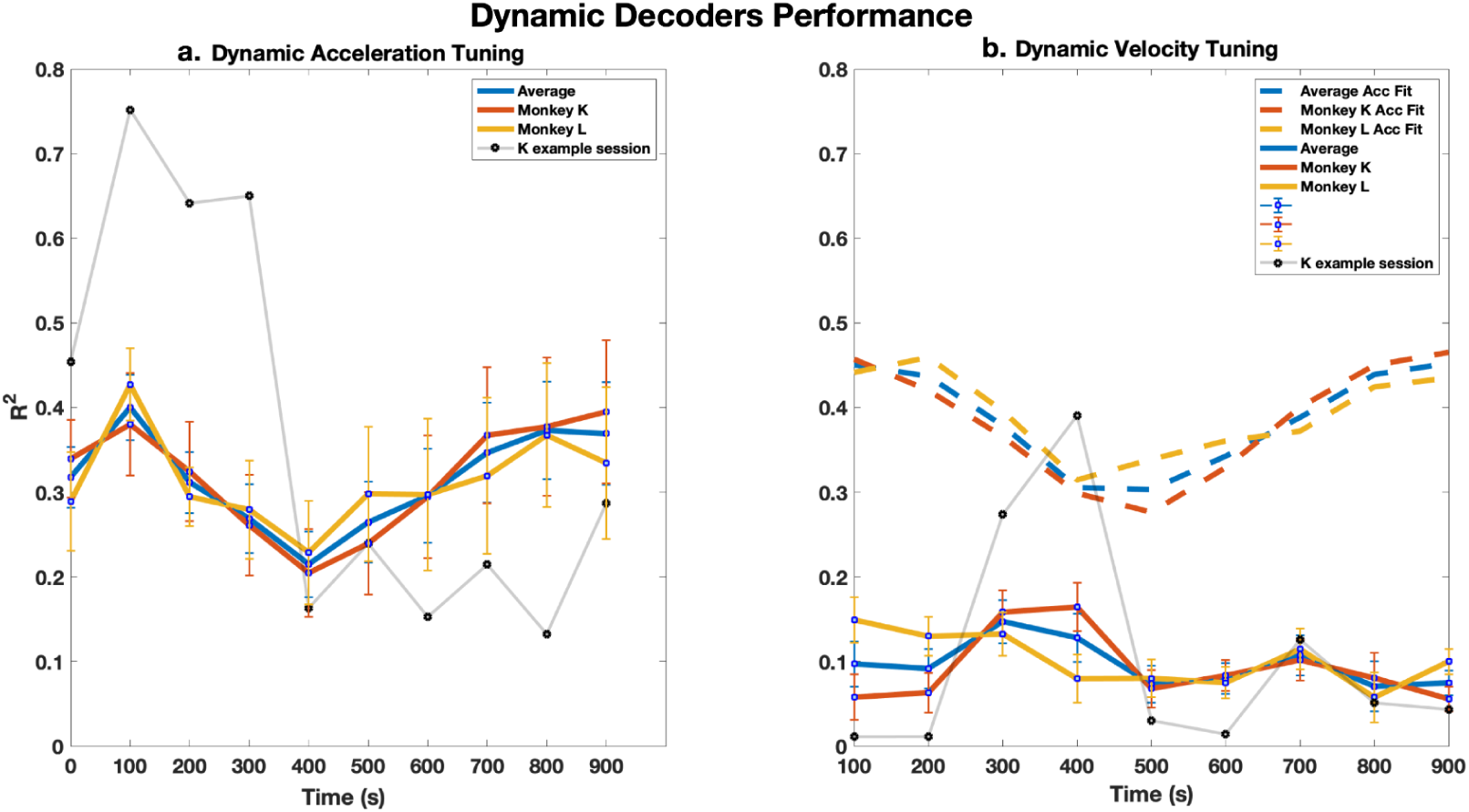
Average Dynamic direct-acceleration decoder performance vs Dynamic velocity-based decoder performance over time. a, Average decoding performance over time of dynamic direct-acceleration decoder for Average (blue line), Monkey K (red line), and Monkey L (yellow line) with error bar of +/- 1 SEM. The light grey line shows the direct-acceleration decoding performance of Monkey K example session. b, Average decoding performance over time of dynamic indirect-velocity decoder (solid lines) with fitted-version of dynamic direct-acceleration decoder performance overlaid (dashed lines). The light grey line shows the indirect-velocity decoding performance of the same Monkey K example session.

**Figure 7.**
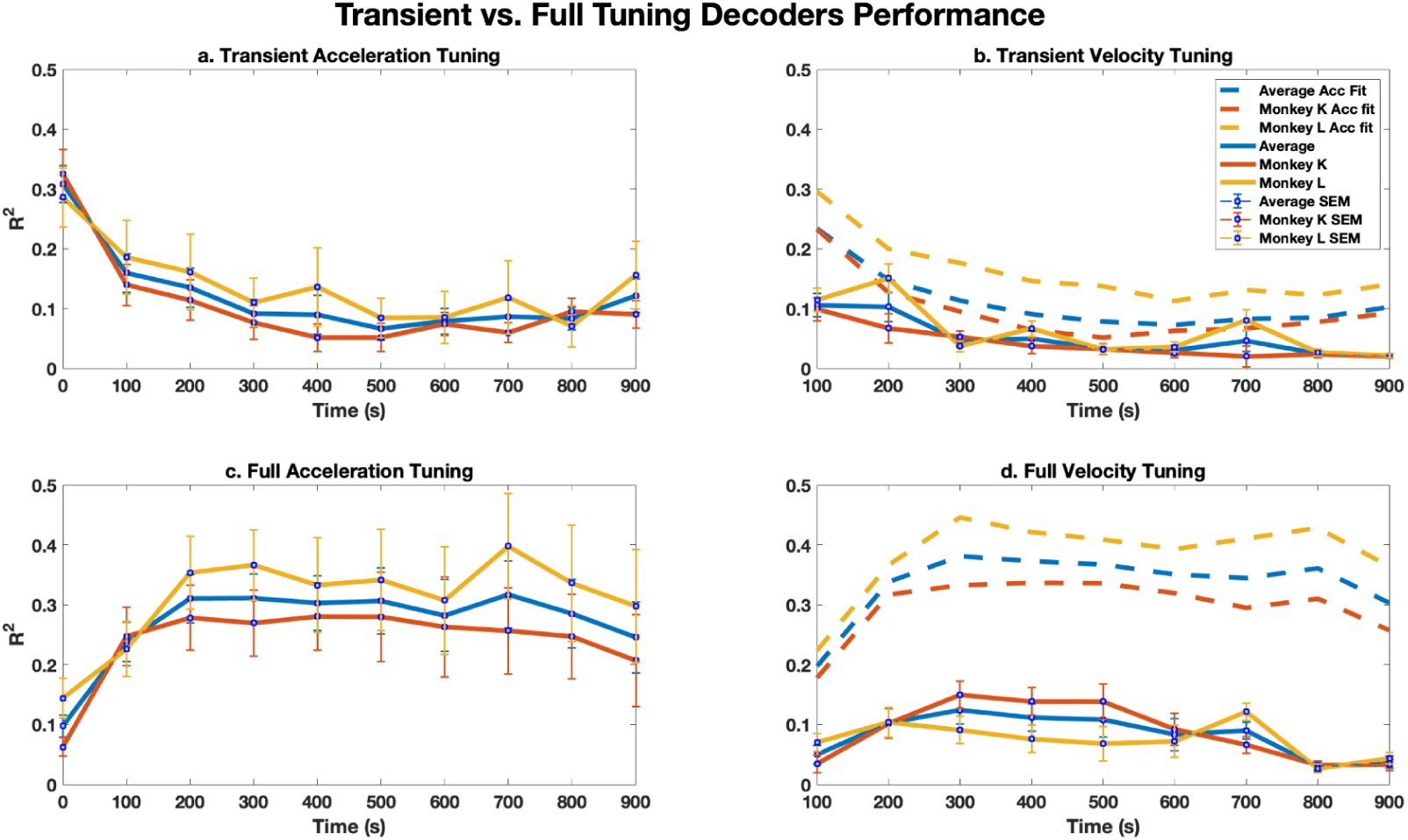
Average transient-tuning and Full-tuning versions of direct-acceleration and indirect-velocity-based decoders performance over time. a, Average decoding performance over time of transient tuning direct-acceleration decoder. b, Average decoding performance over time of transient tuning indirect-velocity decoder (solid lines) with fitted-version of transient tuning direct-acceleration decoder performance overlaid (dashed lines). c, Average decoding performance over time of full-tuning direct-acceleration decoder. d, Average decoding performance over time of full-tuning indirect-velocity decoder (solid lines) with fitted-version of full-tuning direct-acceleration decoder performance overlaid (dashed lines). Average (blue line), Monkey K (red line), and Monkey L (yellow line) with error bar of +/- 1 SEM.

The results from dynamic direct-acceleration decoder reveal the dynamics of MT acceleration readout: the decoding performance started high during the very first 200ms that included the onset transient period, and the performance gradually decreased and then generally increased during the sustained response time (Figure 6a, Dynamic acceleration tuning).

Seeing the temporally-dynamic patterns of the decoding performance, we then further investigated if such dynamics are needed by MT ensemble to efficiently determine motion acceleration condition, and if MT potentially applies different mechanisms to encode acceleration information during transient and sustained response phases during ongoing visual motion. Based on the assumption that the motion onset-transient responses are sufficient to capture the motion acceleration, a “transient” direct-acceleration decoder did (unsurprisingly) decode acceleration effectively in the initial epoch, but this high performance was not maintained throughout the stimulus presentation. The decoding performances in the later epochs decreased and then plateaued at a level much lower than those of the dynamic decoder (Figure 7a, Transient acceleration tuning), indicating that the sustained period needs its own set of decoding weights that are different from the weights utilized by the transient period. What’s more, the average performance results showed that the transient epoch performance drops dramatically compared to the dynamic version, but the sustained epochs performance is maintained at relatively high level without much variance (Figure 7c, Full acceleration tuning).

Applying full-trial acceleration tuning as decoding weights generally gave the sustained period high and stable decoding performance which was sometimes even higher than that from the dynamic version at the same time points and thus indicating the sustained period epochs may share a common decoding mechanism. However, this made it very difficult to accurately and rapidly extract acceleration information within the first 100-200ms after motion onset.

Finally, we looked at the velocity-based counterparts of dynamic (Figure 6b), transient (Figure 7b) and full-tuning (Figure 7d) variations. The dynamic velocity-based decoder seemed to present a similar decoding performance pattern to the direct-acceleration counterpart showing that the decoding strength increased and then decreased over time. However, the decoding strength was generally much lower than that from the dynamic direct-acceleration decoder. The transient velocity-based decoder showed a similar decoding performance pattern to the transient direct-acceleration decoder with the transient interval (first 200ms) having the best performance and sustained intervals with performance dropped close to zero (Figure 7b). In regards to the full-tuning velocity-based decoder results (Figure 7d), the performance increased from near-zero level at the transient interval and then gradually decreased to near-zero level. In sum, velocity-based decoders results showed similar performance changes/variation patterns to their direct-acceleration counterparts across all three variations, but with much lower decoding performance at all time points. To allow for a clearer visualization and more fair and direct comparison to the velocity-based decoders’ performance, we also took the fitted-line version of the direct-acceleration decoders’ estimates. We simply fitted a line between each adjacent pair of epochs and took the mid-value (equivalent to the mean) as the fitted acceleration estimate. The fits are plotted as dashed-lines overlaid on velocity decoders’ results in Figure 6b, Figure 7b and 7d. The same decoding performance patterns were seen across all versions of direct-acceleration decoder.

In closing, the acceleration decoding performance results from variations of direct-acceleration decoders and velocity-based decoders, elucidated that extraction of motion acceleration can be done from MT population responses on a rather fast sub-trial timescale, instead of indirectly estimating acceleration condition from instantaneous changes of velocity during the stimulus presentation. This direct decoding also considers temporal dynamics of the MT neural activity throughout the ongoing visual motion, so the neural population is actively and continuously pooling and weighing the encoded acceleration information represented in its units’ firing. Specifically, the brain may employ different decoding strategies or mechanisms with motion onset transient responses and sustained responses respectively to first efficiently decode acceleration at the very start of the visual motion and then maintain a relatively high level of performance till the motion offset.

### Area MST does not refine acceleration representations

Area MST, which receives massive direct input from area MT, is usually considered as a higher and later stage of visual motion processing that prefers complex motion patterns (Celebrini & Newsome, 1994; Huk et al., 2002; Raiguel et al., 1997; Wild & Treue, 2021). With the evidence that we can directly and rapidly decode motion acceleration from MT ensemble activity, we further asked if MST could refine this acceleration representation. To achieve this, we recorded from the same two monkeys’ MST ensembles in separate sessions with the same experimental setup (Figure 8) and applied the same analysis for decoding performance (Figure 9).

**Figure 8.**
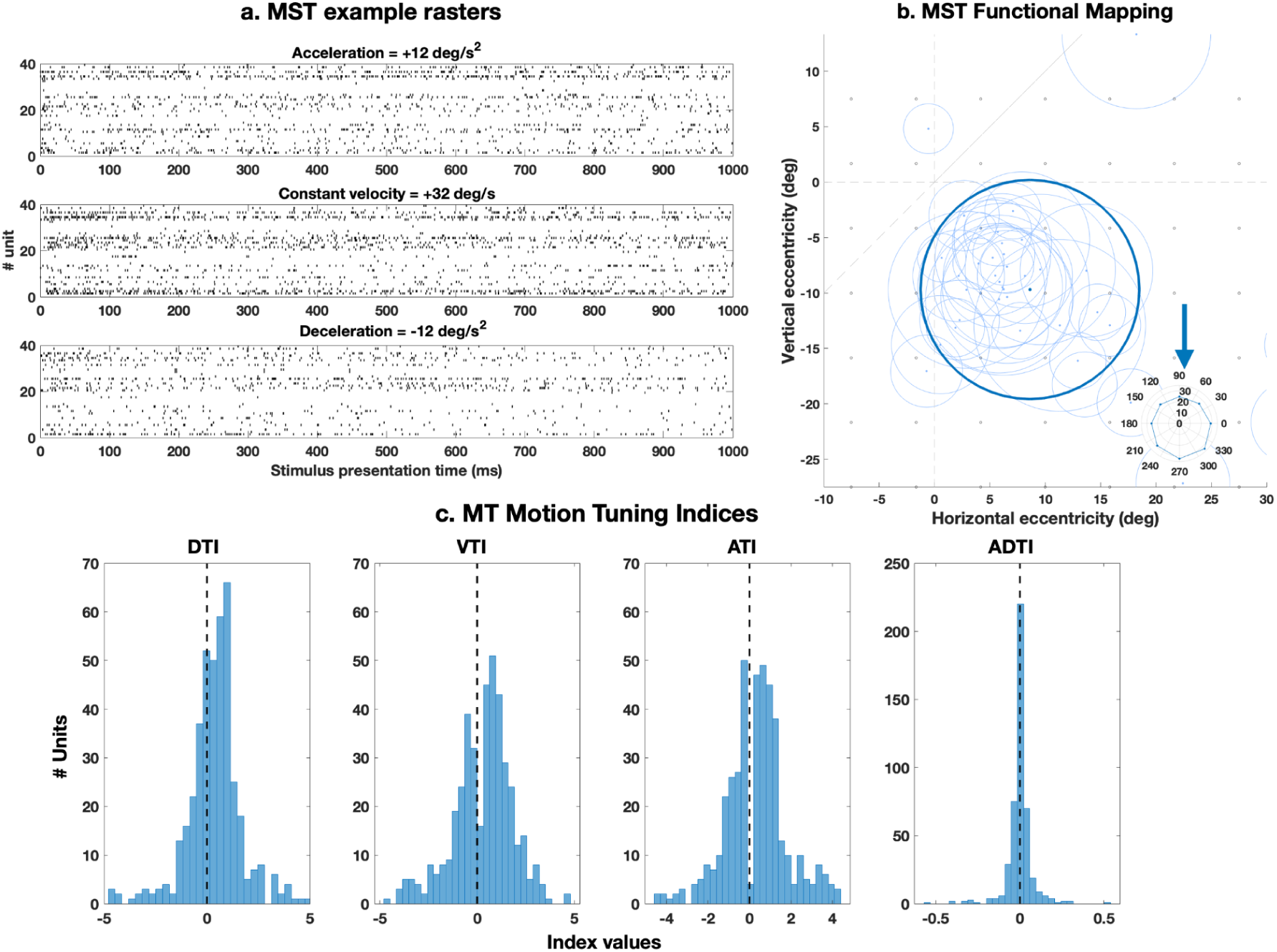
MST functional mapping. a, MST example trial rasters for acceleration (top), constant velocity (middle) and deceleration (bottom). b, MST functional mapping: light blue contours show individual RFs and the bold contour indicates the location and size for the acceleration stimulus envelope. The blue arrow shows the direction of preferred-null motion axis with the inset polar plot showing the population direction tuning. c, Histograms of motion tuning index from MST sessions, left–right: Direction tuning index (t(460)=5.632, p<0.001, M=0.366, SD=1.234), Velocity tuning index (t(460)=4.180, p<0.001, M=0.308, SD=1.509), Acceleration tuning index (t(460)=4.538, p<0.001, M=0.329, SD=1.482) and Acceleration-deceleration tuning index (t(460)=-0.210, p=0.834).

**Figure 9.**
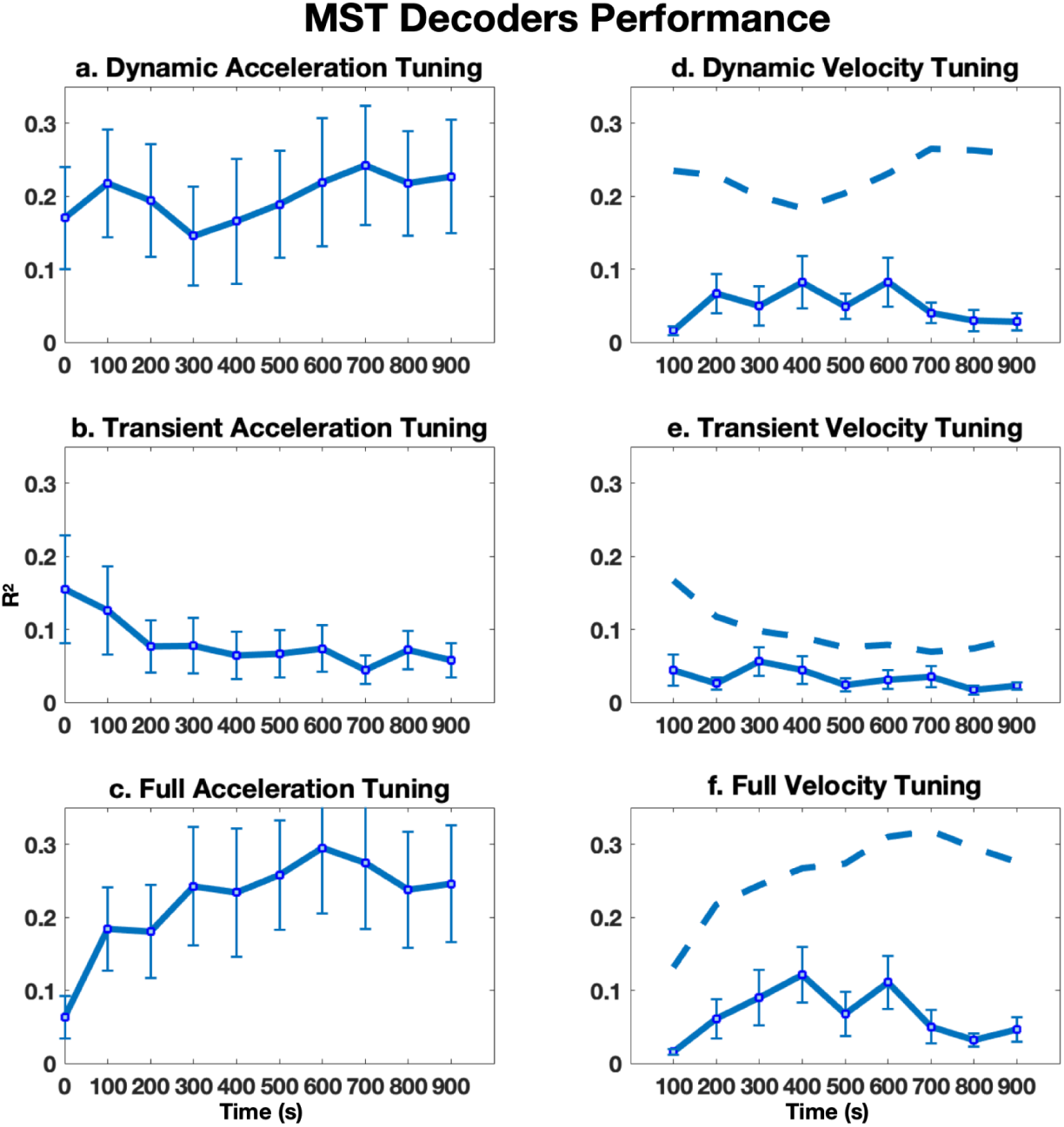
MST acceleration decoding decoders performance. Average decoding performance over time of **a**, Dynamic tuning direct-acceleration decoder. **b**, Transient tuning direct-acceleration decoder. **c**, Full-tuning direct-acceleration decoder. **d**, Dynamic tuning indirect-velocity based decoder (solid line). **e**, Transient tuning indirect-velocity based decoder. and **f**, Full-tuning indirect-velocity decoder. Dashed lines are the linear fitted version of direct-acceleration decoder results for comparison to velocity-based decoders. Error bars are +/- 1 SEM.

Figure 8 c’s histograms summarize the visual motion tuning strength of MST. Explicitly, 59.22% (n = 273) of the recorded MST single units showed preferred acceleration with 31.67% (n = 146) excited by it and 40.78% (n = 188) were inhibited by all acceleration conditions.

Similarly, 59.87% (n = 276) of the units are velocity tuned with 34.92% (n = 161) excited by the preferred velocity and 40.13% (n = 185) were inhibited by any visual motion velocity. As for direction and binocular disparity tuning, 60.52% (n = 279) of the neurons preferred specific motion direction or disparity, but 38.39% (n = 177) were inhibited by all sampled directions and disparity levels. Within the tuned units, 29.50% (n = 136) and 32.54% (n= 150) of them were excited by their preferred direction or disparity. In general, the acceleration, velocity, direction and binocular disparity tuning strengths of the recorded MST units were lower than those of the MT units for both monkeys and on average. Moreover, compared to MT units, MST units showed more even preference over positive accelerations and negative accelerations (decelerations) as 49.89% (n = 230) of them prefer acceleration, 47.94% (n = 221) prefer deceleration and the rest 2.17% (n = 10) prefer constant velocity (MT ADTI and MST ADTI distributions are significantly different: t(648.233)=3.938, *p*=0.021, M=[0.021, <0.001], SD=[0.108, 0.079]). Though in general MST units showed lower tuning strength compared to MT units, the single units in MST did show comparable tuning across motion dimensions similar to MT’s motion tuning (DTI: t(358.157)=-1.065, *p*=0.288, M=[0.857, 0.366], SD=[1.070, 2.154]; VTI: t(557.922)=0.120, *p*=0.905, M=[0.776, 0.308], SD=[1.470, 2.995]; ATI: t(459.238)=0.459, *p*=0.646, M=[0.658, 0.328], SD=[1.362, 5.132]).

Next, similar to the MT results, we checked on the decoding strength of versions of MST direct-acceleration and velocity-based decoders (Figure 9). For dynamic tuning decoders, akin to what have been seen with MT results, the decoding strength was high at the start and then had decrement and increment followed for direct-acceleration decoder, and in contrast the decoding performance was generally low and close to zero level across time for velocity-based decoder (Figure 9 a&b). Though MST in general preserves the direct-acceleration decoding temporal dynamics, the strength of decoding was much lower across the stimulus presentation time. The epochs that cover the initial 200ms of the transient direct-acceleration decoder did show slightly higher decoding performance with other epochs having near zero performance (Figure 9b). The transient velocity-based decoder basically showed almost no decodability to motion acceleration (Figure 9e). As for the full-tuning version of MST acceleration decoders, on average both direct-acceleration decoder and velocity-based decoder exhibited comparable levels of decoding strength and patterns to those results from MT (Figure 9c&f). With the adjusted-R^2^ values derived as the representation of decoding strength for each epoch or interval of the trial, we then looked at the distributions of these values of dynamic direct-acceleration decoder and velocity-based decoder for both MT and MST. Mostly, the R^2^ values of dynamic direct-acceleration decoder from both MT (mean = 0.3163) and MST (mean = 0.1978) were remarkably higher than those from dynamic velocity-based decoder (MT mean = 0.0967, MST mean = 0.0493) (Figure 10). This further supported the idea that motion sensitive areas applying direct and dynamic acceleration decoding is more efficient and accurate than making inference about acceleration from velocities. In summary, acceleration is directly decodable from the MST ensemble with its temporal dynamics, and potentially a different decoding mechanism for motion onset from the rest of the motion is applied. However, MST does not seem to be refining acceleration decoding or even maintaining the high decoding performance seen in MT ensembles. In addition, MST does not seem to make use of velocity-based decoding as an alternative to indirectly extract motion acceleration.

**Figure 10.**
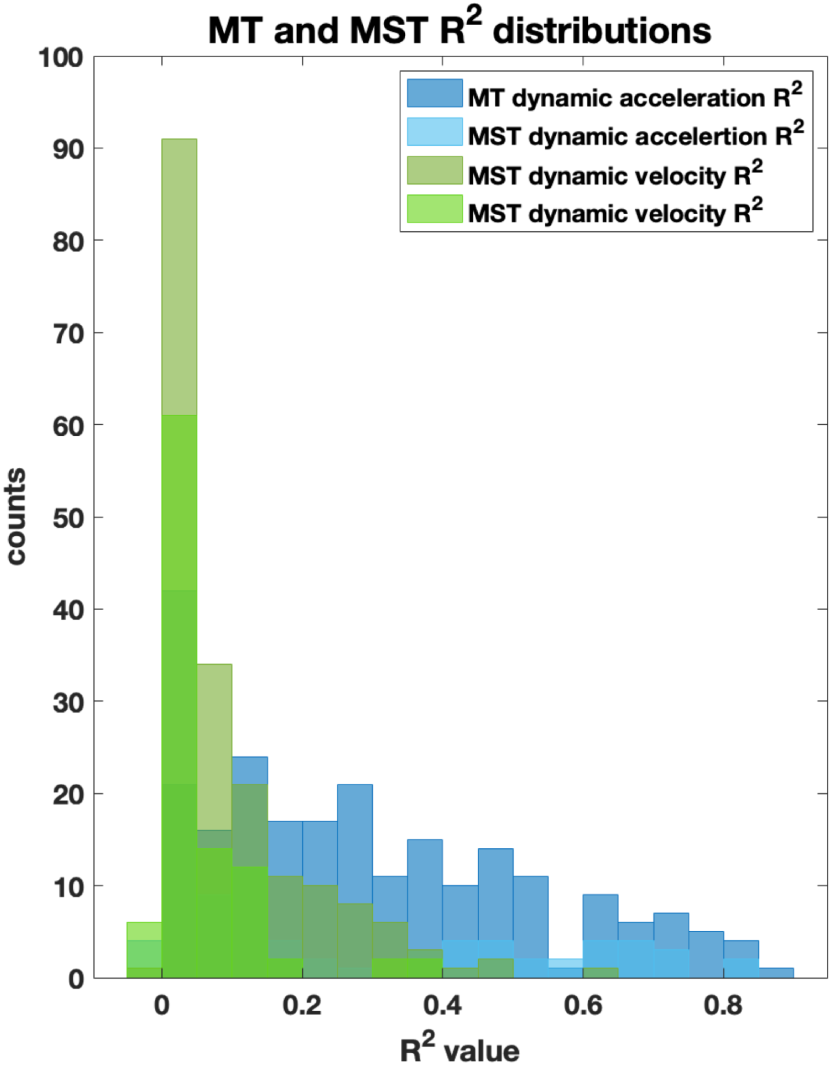
MT and MST dynamic acceleration and dynamic velocity decoders’ R^2^ distributions. Blue histograms show the R^2^ distributions of dynamic direct-acceleration decoders, and the green histograms show the R^2^ distribution of dynamic indirect-velocity based decoders.

## Discussion

We studied both the single-neuron and population-level representations of visual motion acceleration in area MT of the macaque. Although this topic has been investigated before in detailed and thoughtful work by Lisberger, Movshon, Price, and others (Cao et al., 2004; Lisberger & Movshon, 1999; Price et al., 2005), the preceding work laid out ideas that we realized could be more definitively addressed with the application of newer techniques both for neural recording and for data analysis. We used three novel approaches to build off that prior work. First, we used multi-electrode arrays to record from ensembles of neurons, which provided us both a more comprehensive sample of MT neurons, as well as performing exploratory recordings in MST more feasible. Second, we sampled a range of accelerations that were limited to fall within perceptually-relevant ranges, which may have helped us characterize tuning in a parameter space that is large and less well-charted than other parameters such as speed or temporal frequency. Third, we performed population decoding analyses that leverage neural responses in a mechanistically-agnostic manner, which could provide better statistical power while sidestepping issues of how such responses come to be and/or whether they fit within distinct information-processing channels. In this Discussion, we first explain in greater detail how our updated techniques provided insights that are consistent with prior work, but which provide more definitive insight into how the brain infers acceleration. We then discuss some new questions that arise from the application of these tools, for the sake of motivating future work.

Perhaps our most novel result is the simple discovery of clear tuning for acceleration at the single-unit level in MT. Although the take-home from prior work might be put simply as a lack of single-unit acceleration tuning, we believe our results are in fact not inconsistent with the results of prior studies. First, our larger samples and perceptually-relevant stimulus range likely contributed to our ability to find neurons with responses whose responses changed systematically across changes in acceleration. Smaller-sample recording techniques may not just miss neurons with tuning, but could also engender biased sampling of neurons with robust responses to other properties that are not predictive of acceleration tuning. Second, the criteria for defining a neuron as tuned are somewhat different across studies (Butts & Goldman, 2006; Kriegeskorte & Wei, 2021). In prior work, there was a strong emphasis on identifying acceleration tuning distinct from other (known) mechanisms, such as velocity tuning and short term adaptation (Price et al., 2005, 2006; Schlack et al., 2007); with time-varying stimuli, this is indeed a thorny problem. We took a more modern population-centric perspective, asserting that behavior only needs access to neural responses that are informative about acceleration, regardless of the mechanistic source of that information (as well as whether the tuning is explicit at the level of single neurons). This way of thinking motivated our treatment of conditions (in which we separated trials by acceleration, irrespective of the starting or ending velocity). It is possible that perceptual estimates of acceleration may be affected by starting or ending velocity, but our analyses show that MT neurons exhibit clear tuning even in the face of variation in those related parameters. In addition, we see the acceleration tuning at both the single cell and ensemble level changed over time, especially with differences between initial, onset-transient tuning and later, sustained period tuning. This indicates that MT’s acceleration encoding may already provide dynamic information for downstream areas to employ different decoding strategies (e.g., transient response decoding vs. sustained response decoding) to both efficiently and accurately extract current motion acceleration conditions.

Our use of now-standard decoding analyses also extends our understanding of acceleration representations in the primate cortex (Berens et al., 2012; Chaisanguanthum & Lisberger, 2011; Chen et al., 2015; Graf et al., 2011; Tanabe, 2013). In the preceding work that strongly motivated this study, Lisberger and Movshon (1999) pointed out that “acceleration could be reconstructed if each neuron’s output was weighted according to the product of preferred speed and a measure of the size of its transient response” (we note that stimulus “reconstruction” was that era’s parlance for “decoding”). In that work, they analyzed the responses of ∼100 individually-recorded MT units and used a model-based framework to perform decoding of acceleration by taking the responses of speed-tuned neurons and weighting them by a term associated with each unit’s degree of transient-to-sustained firing. This model thus combined information about speed— uncontroversially encoded in MT units— with each neuron’s particular temporal dynamics that could be used to infer the timing of a particular speed being encoded.

Our approach differs from these earlier reconstruction/decoding approaches by sidestepping prior foci on the presence of well-defined signal channels or interpretable response components (such as relying on sustained vs. transient responses, or parcelling out out effects of adaptation), and instead explores the notion of “opportunistic decoding”, in which idiosyncrasies of neural responses can be exploited. In the specific case of acceleration, the temporal dynamics of responses not simply explainable in terms of linear stimulus filtering reflect nonlinearities in the temporal domain. Although nonlinear interactions between stimulus and task dimensions are now widely recognized to provide computational advantages for decoding (Albrecht et al., 2003; Cohen & Newsome, 2008; Orhan & Ma, 2015; Yang et al., 2021; Yiling et al., 2024), our work here highlights a related, but less appreciated, insight: nonlinear interactions between a stimulus/task variable and time itself are also useful for decoding (Botella-Soler et al., 2018; Hajnal et al., 2023; Tajima et al., 2017). In our analyses, they allow the brain to infer a stimulus property more quickly than would be possible with a temporal cascade of linear operations. Because fast timescale adaptation is a widespread temporal nonlinearity in the brain (Bair & Movshon, 2004; Dragoi et al., 2000; Kohn & Movshon, 2003, 2004; Patterson et al., 2014; Price & Born, 2013; Priebe & Lisberger, 2002; Teich & Qian, 2003), it is possible that our observations in this specific domain reveal a general “trick” for decoding on fast time scales.

This issue leads our discussion to the first of a few new questions that are brought into focus by our study. We do not know if the brain needs well-defined neural channels or temporal response phases to facilitate decoding of stimulus properties, or whether the brain’s read-out mechanisms can more agnostically exploit neuronal selectivity and idiosyncrasy with the same adeptness as the statistical exercise implemented via our simple population decoder. It will be intriguing to see future work attempt to understand how the brain flexibly weights population activity to flexibly and rapidly perform such read outs. The use of tools to perturb certain cell types and projection pathways make this endeavor now seem viable, even in some primate species (Klein et al., 2016; Li & Liu, 2023; Merlin & Vidyasagar, 2023; Oguchi & Sakagami, 2022; Upright et al., 2018).

Another unanswered question brought into focus by our study is that the representation of acceleration did not appear to be further refined in area MST. MST is well-known to exhibit tuning for complex spatiotemporal patterns of motion likely related to optic flow processing (Beardsley & Vaina, n.d.; Graziano et al., 1994; Mineault et al., 2012; Paolini et al., 2000; Yu et al., 2010). We believed that its status as a “later” stage of motion processing warranted exploration to see if it took velocity signals from MT and further calculated acceleration from those inputs. We did not find evidence for this. Instead, our MT decoding results suggest that an explicit, serial calculation of temporal derivatives is not needed and that acceleration information is available via fast and direct decoding based on temporal nonlinearities.

Finally, our finding of single-unit tuning for acceleration is novel and likely resulted from a fortuitous choice of stimulus parameters (i.e., a useful range of accelerations and sampling along that range with the resolution to identify systematic tuning). Other stimulus parameters (not directly related to acceleration) may affect MT responses, and whether a decoding scheme needs to flexibly adjust to handle such “nuisance parameters” is an important topic for further work. Analyses of decoding quality and flexibility would be best guided by matched psychophysical studies to assess how well real brains are able to infer acceleration in the face of other parameters that affect neural signalling but do not provide additional information about the variable of interest.

## Acknowledgements

This work was supported by NEI R01 EY020592. We thank Thaddeus Czuba and Lawrence C Cormack for assistance with the design and programming of visual stimuli. We also thank Allison Laudano and Declan Rowley for assistance with animal surgery, care and training.

